# Fecal microbiota transplantation brings about bacterial strain displacement in patients with inflammatory bowel diseases

**DOI:** 10.1101/439687

**Authors:** Manli Zou, Zhuye Jie, Bota Cui, Honggang Wang, Qiang Feng, Yuanqiang Zou, Xiuqing Zhang, Huanming Yang, Jian Wang, Faming Zhang, Huijue Jia

## Abstract

Fecal microbiota transplantation (FMT), which is thought to have the potential to correct dysbiosis of gut microbiota, has recently been used to treat inflammatory bowel disease (IBD). To elucidate the extent and principles of microbiota engraftment in IBD patients after FMT treatment, we conducted an interventional prospective cohort study. The cohort included two categories of patients: (1) patients with moderate to severe Crohn’s disease (CD) (Harvey-Bradshaw Index ≥ 7, n = 11, and (2) patients with ulcerative colitis (UC) (Montreal classification, S2 and S3, n = 4). All patients were treated with a single FMT (via mid-gut, from healthy donors) and follow-up visits were performed at baseline, 3 days, one week, and one month after FMT (missing time points included). At each follow-up time point, fecal samples of the participants were collected along with their clinical metadata. For comparative analysis, 10 fecal samples from 10 healthy people were included to represent the diversity level of normal gut microbiota. Additionally, the metagenomic data of 25 fecal samples from 5 individuals with metabolic syndrome who underwent autologous FMT treatment were downloaded from a previous published paper to represent natural microbiota shifts during FMT. All fecal samples underwent shotgun metagenomic sequencing.

We found that 3 days after FMT, 11 out of 15 recipients were in remission (3 out of 4 UC recipients; 8 out of 11 CD recipients). Generally, bacterial colonization was observed to be lower in CD recipients than in UC recipients at both species and strain levels. Furthermore, across species, different strains displayed disease-specific displacement advantages under two-disease status. Finally, most post-FMT species (> 80%) could be properly predicted (AUC > 85%) using a random forest classification model, with the gut microbiota composition and clinical parameters of pre-FMT recipients acting as the most contributive factors for prediction accuracy.

## INTRODUCTION

Inflammatory bowel disease (IBD) is a chronic inflammatory disease characterized by chronic immune-mediated intestinal inflammation, and consists mainly of Crohn’s disease (CD) and ulcerative colitis (UC). The etiology of IBD has been proposed to be multifactorial, involving a dysregulated immune response to environmental factors in a genetically susceptible individual (*1*). Interestingly, given the evidence accumulated in recent years, the gut microbiota is now recognized for playing an important role in IBD. Dysbiosis is a decrease in gut microbial diversity owing to a shift in the balance between commensal and potentially pathogenic microorganisms of the gut microbial ecosystem, and has long been characterized as a trait of IBD patients (*2,3*).

Fecal microbiota transplantation (FMT) aims to modify the intestinal microbiota composition and function of the recipients by transferring donor fecal suspension into the gastrointestinal tract of a recipient, and has become a promising method for manipulating the gut microbiota. Its successful application for the treatment of *Clostridium difficile* infection has inspired people to apply it to inflammatory bowel disease patients (*4,5,6,7,8,9*). However, this application is still in its early stages. According to a recent systematic review and meta-analysis, after minimizing publication bias, IBD patients who received FMT had a remission rate of only 36.2%: 22% for UC and 60.5% for CD (*10*). Moreover, there is a lack of research regarding the efficiency and principles of FMT in treating IBD.

Clinical research to date has focused more on UC (*7,8,9*), and there has been insufficient research on the effects of FMT on CD patients, with only a few case reports and small-scale case series reported (*11,12,13,14*). In addition, the majority of studies conducted so far to investigate the role FMT plays in treating IBD have used 16S rRNA sequencing, which has limited resolution on taxonomic and functional classification of sequences. Contradictory results were often observed at species-level resolution, making it hard to determine the exact role of different bacterial agents. For instance, the abundance of *Faecalibacterium prausnitzii* was found to decrease in one study and to increase in another (*15,16*). Thus, it is necessary to be able to appreciate the whole composition of gut microbiota at a strain level. Strain level variants within microbial species are crucial in determining their functional capacities within the human microbiome, such as interaction with host tissues (*17*), modulation of immune homeostasis (*18*), and xenobiotic metabolism (*19*). Shotgun metagenomic sequencing with the ability to target all DNA material in a sample can give a base pair level resolution of the genome that makes single nucleotide analysis possible. Additionally, promising machine learning methods could enable the establishment of predictive models to predict the microbiota composition of post-FMT recipients. Recently, *S. Smillie* et al. constructed a machine learning model to predict the species profile of post-FMT recipients for 18 *C. difficile* patients and found that bacterial abundance and phylogeny were the strongest determinants of engraftment (*20*). In our study, we utilize a random forest model to predict the mOTUs profile of IBD recipient 3 days after FMT and identified the variables that contribute most to model prediction accuracy.

## MATERIALS AND METHODS

### Patient recruitment and sample collection

Patients aged 19–64 years were recruited from the Second Affiliated Hospital of Nanjing Medical University, China from 2012 to 2014. The dataset was composed of 10 fecal samples from 10 healthy people, among which 6 were FMT donors, and 34 fecal samples from 15 IBD patients. Donor fecal samples were collected prior to FMT. Stool samples from recipients were collected at baseline, day 3, and day 7 (or day 30) (Figure 1). Missing points were due to patient discharge. Detailed standards of patient recruitment and donor screening were previously published (*13*). Donors were either related (genetically related family members) or unrelated (screened unrelated family members). Clinical metadata of IBD patients—including anthropometric index, clinical parameters, and blood test results—were obtained at each follow-up time point. For autologous FMT treatment, 25 additional fecal samples from 5 metabolic syndrome individuals were obtained from the *Vrieze* et al. (21) study with follow-up points on day 0 and days 2, 14, 42, and 84 after FMT.

**Figure 1.**
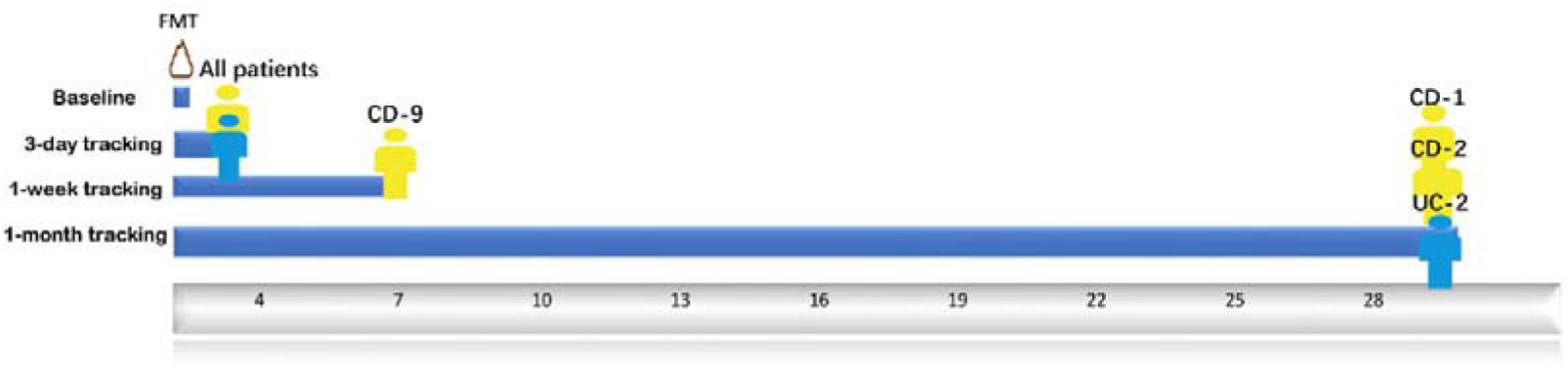
**Study design and follow-up visits of the patients** To recognize each patient in simplicity, we labeled each of them with disease subtypes CD- or UC- as prefix plus a random assigned number as suffix.

In summary, 34 samples were used for analysis of the allogenic FMT group, 25 for the autologous, and 10 for the healthy group.

### Stool sample collection and FMT procedure

Fecal samples were obtained from scanned donors and were isolated for microbiota at lab. Fecal microbiota from the donor was prepared according to the manual method of filtration, centrifugation, washing, discarding, and resuspension and repeated processes. Purified fresh fecal microbiota suspension was input into patients’ mid-gut by a tube within gastroscope under anesthesia, and the entire procedure should be done within one hour.

### Metagenomic sequencing and processing methods

DNA extraction and metagenomic sequencing of IBD fecal samples and healthy fecal samples were performed at BGI-Shenzhen, China following HiSeq 2000 sequencing protocol. Metagenomic sequencing of autologous FMT treatment samples was performed at the Genomics Core Facility of the European Molecular Biology Laboratory, Heidelberg using Hiseq 2000.

Illumina sequencing reads were quality controlled by trimming low quality bases (quality score < 20), filtering adapter reads, and removing host-related reads after mapping to the human genome database. The reads quality control procedure was conducted using cOMG with default parameters (*22*). After quality control, 1,379,430,125 sequences were obtained, with a mean of 31,350,685 sequences per sample.

### Microbiota taxonomic profiling

Species-level quantification of metagenomic sequencing reads was achieved using mOTUs software with default parameters. mOTUs is a method that establishes metagenomic operational taxonomic units based on single-copy phylogenetic marker genes. It maps the quality-controlled metagenomic sequencing reads against the m-OTUS.v1.padded database, which is composed of 10 MGs extracted from 3,496 prokaryotic reference genomes (download from NCBI) and 263 publicly available metagenomes (from the MetaHIT and HMP projects), and then outputs metagenomic OUT linkage groups (m-OTUS) (*23*).

For strain level profiling, metaSNV was utilized to process quality-controlled metagenomic sequencing reads. metaSNV is a method that is able to disentangle conspecific strains in metagenomic samples using specific single-site allelic variation (SNVs). It uses a collection of microbial reference genomes in which each species is represented by a single representative genome or gene collection (*24*). To maintain consistency with previous species profiles, we specified the m-OTUS.v1.padded database as our reference genome or gene collection during this procedure. First, we mapped quality-controlled sequencing reads to the m-OTUS.v1.padded database using bwa and Ngless. Next, we ran qaCompute on each sample to determine the average coverage over each reference in each sample and aggregated the coverage information. We then took advantage of the mpileup tool to compute genomic variation, and outputted all the variant positions that met the default-imposed quality criteria. Lastly, we computed per species pairwise distance matrices for the samples.

### Quantification and Statistical Analysis

All statistical analyses were performed in R using the following packages: vegan, Hmcc, pROC, and RandomForest. We conservatively used only the baseline and day 3 time point samples for each patient when conducting all the two-sided statistical tests.

#### Diversity comparisons

The diversity of each gut microbiota community per sample was calculated based on its mOTUs profile, referred to as the Shannon index, using the vegan package. The Kruskal-Wallis test was used as a significance test for this multi-group comparison.

#### Species-level changes after FMT

After species profiling all fecal samples using mOTU, we took only the species with a detected relative abundance of at least 0.001 into account to avoid ambiguous results. In order to determine whether donor microbiota could be transferred to recipients, we divided the microbiota composition of post-FMT recipient into 4 groups: donor-specific species, recipient-specific species, common species (shared by donor and recipient), and new species (not found in either the donor or in the pre-FMT recipient). We quantified these 4 groups by comparing the gut microbiota mOTU profiles of the pre-FMT recipient, the post-FMT recipient, and the donor. Results were visualized using bar plota with all available follow-up time points.

#### Community-level changes after FMT

Community-level changes in gut microbiota composition between pre-FMT and post-FMT recipients were represented by the Bray-Curtis distance, which was computed using the vegan package after applying a logarithmic transformation to mOTU relative abundance with the function log(x+x_0_), where x is the original relative abundance of a certain mOTU and x_0_ = 1e-6. The cosine dissimilarity was also used to examine the correlations between gut microbiota compositions pre-FMT and post-FMT, and between post-FMT recipients and donors. Results were displayed using scatter plots.

#### Strain-level changes after FMT

Strain differentiation, which was determined by comparing the presence or absence of donor-specific, recipient-specific, and previously undetected single-site allelic variations, was monitored in post-FMT recipients based on the output files of metaSNV. Similar to the process of determining species retention and transplantation, the gut microbiota composition of post-FMT recipients was categorized into 3 groups: donor-specific strains, recipient-specific strains, and common strains (shared by donor and recipient). We excluded the newly gained strains because that was not of interest here. Quantification of the three groups was determined according to the frequency per filtered SNVs set.

#### Species engraftment model

We sought to investigate whether the microbiota composition of post-FMT recipients could be predicted using advanced machine learning models. We therefore applied the Random Forest algorithm in R to predict the presence (random forest classification model) and abundance (random forest regression model) of each mOTU in every post-FMT recipient sample. For a dataset comprised of 15 samples and 123 filtered mOTUs, these models are trained on 15 x 127 total instances. The inputs for these predictions are the gut microbiota composition of each pre-FMT patient and their corresponding donor at a species level, along with clinical metadata of the pre-FMT recipient and donor. Random Forest is a collection or ensemble of classification and regression trees trained on targeted datasets. It is resistant to overfitting and is considered stable in the presence of outliers. The error rate of the classification of all the test sets is the out-of-bag (OOB) estimate of the generalization error (*25*).

First, we eliminated the condition of class imbalances by filtering out mOTUs that existed in less than 3 samples to avoid prediction bias in favor of the majority class.

Second, the mtry parameter with the lowest error was picked using the rfcv function with 5-fold cross validation. Third, we applied the randomForest function to perform classification of post-FMT recipients across all mOTUs. This resulted in 123 randomForest classification models in total, and we computed the auc value for each model. Finally, we chose important features from those models that had good prediction performance (auc bigger than 0.9).

For the regression model, we also accounted for class balance and then used the rfcv function with the same predictors that we used in the classification model to perform prediction.

#### Feature Importance

Random Forest calculates feature importance by removing each feature from the model and measuring the decrease in accuracy (for presence) or the increase in the mean-square error (for abundance). According to these importance scores, we ranked features in decreasing order across models and picked 40 with the highest scores to display.

#### Correlations between change in mOTUs as well as in clinical parameters

Clinical metadata of patients was collected at baseline and follow-up visits, including physical parameters, inflammation markers, lymphocyte population, blood fat, and immunoglobulin. We used the rcorr function in the Hmisc package to compute the spearman correlation iterating from each mOTU-clinical index pair. The change in each mOTU was defined as the increase or decrease in its relative abundance 3 days after FMT treatment compared to baseline. Changes in clinical index were computed based on the absolute score recipients got at baseline and 3 days after FMT treatment.

For multiple comparisons, the Benjamini-Hochberg method was used to adjust the p value to control for false positives. Lastly, we drew a network using Cytoscape based on the pairs with a q-value smaller than 0.05 (*26*).

### Ethical statement

This study was carried out in accordance with the recommendations of good clinical research practice (GCP), the Ethical committee of the Second Affiliated Hospital of Nanjing Medical University, and BGI-IRB. The protocol was approved by the Ethical committee of the Second Affiliated Hospital of Nanjing Medical University and BGI-IRB. All subjects gave written informed consent in accordance with the Declaration of Helsinki.

## RESULTS

### Bacteria characterization at a species level

After profiling sequenced fecal samples using shotgun metagenomics, the Shannon index (alpha diversity of a community) of gut microbiota was measured across IBD recipients. Results showed that the average Shannon index of CD patients was significantly lower than that of healthy controls (P-value = 0.0035). In UC patients, although their Shannon index was lower than the average in healthy controls, dysbiosis was not significant (p-value = 0.57). Three days after FMT treatment, the average Shannon indexes of both CD and UC recipients had not significantly improved (p-value > 0.01) (Figure 2A). Unexpectedly, CD-6, CD-7, CD-8, and UC-2 had a decreased Shannon index.

**Figure 2.**
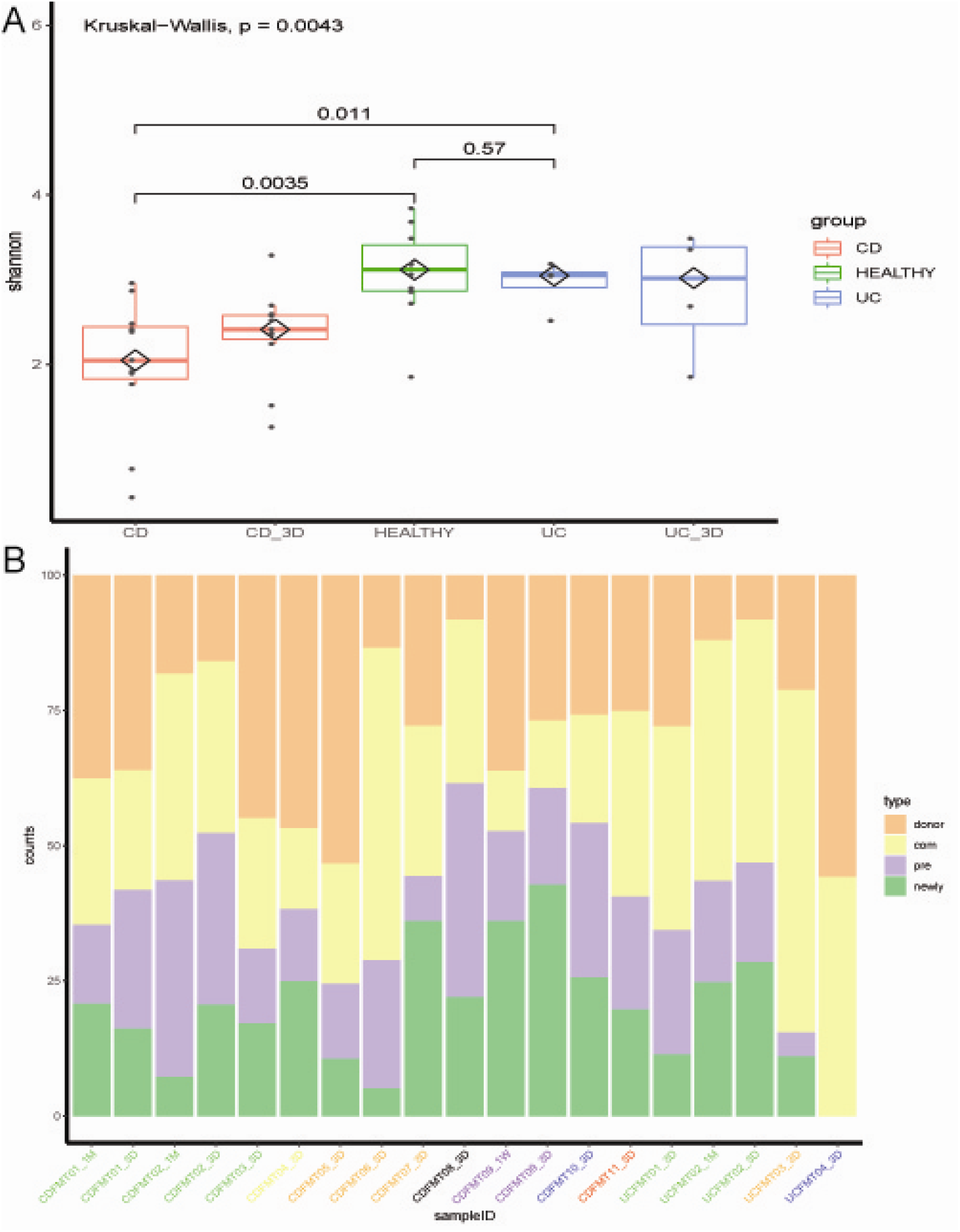
**Bacterial communities undergo compositional changes in IBD recipients after FMT.** (A) The Shannon index of gut microbiota was lower in IBD patients than in healthy controls, and was not significantly improved 3 days after FMT (p-value > 0.01). Different groups are represented by different colored boxes. (B) The proportion of species gained from the donor in post-FMT recipients lasts during follow-up visits. However, the proportions varied among recipients, even those who shared a donor (labels with the same color). Gut microbiota composition per patient was divided into four parts: orange represented donor-specific species, yellow represented species shared by donor and recipient, purple represented recipient-specific species and green represented newly gained species.

Among the whole population of the gut microbiota, some bacteria may be more important than others for maintaining a healthy gut environment. For example, 3 days after FMT treatment, there was a universal increase in *Bacteroides* that have been shown to exist at lower levels in IBD patients than in healthy people (*27*). Some highly individualistic performances were also observed: CD-9 gained an abundant amount of *Lactobacillus*, which was considered to be probiotics, and CD-1 had a great decrease in *Citrobacter*, which was recognized to be pathogenic bacteria (Figure 2B). The amounts of species each recipient gained from their donor after FMT are shown in Figure S1.

### Bacterial engraftment at the species level

To investigate the extent to which the gut microbiota of recipients could be altered by FMT treatment, we evaluated both the degree and direction of change. Results showed that microbial communities underwent large compositional changes after FMT, and these changes persisted throughout follow-up visits (Figure 2B).

On average, post-FMT CD recipients gained 29.4% of mOTUs from donors (n = 11, SD = 14.4%), while post-FMT UC recipients gained 28.2% of mOTUs from donors (n = 4, SD= 20%). Our results were analogous to a previous study that found that FMT recipients gained 35% of mOTUs from donors (n = 436, SD = 27%) (*28*).

By measuring the distance between donor-recipient pairs using Euclidean distance, we determined the direction of microbiota change. Results varied between different donor-recipient pairs. Out of the 4 patients that had 2 follow-up time points, we found that CD-9 and UC-2 tended to be closer to their donors and further from their pre-FMT status. CD-2 showed a slightly tendency to return to their initial status, but the disturbance was small enough to be ignored (a shift from 10.628 to 10.57).

Surprisingly, CD-1 showed an increased distance from both their donor and their pre-FMT status, which could be attributed to environmental factors. Though CD-1, CD-2, and UC-2 all shared the same donor, the direction of their gut flora shift after the treatment varied (Figure 3A). In addition, we explored the abundance consistency of mOTUs of recipients before and after FMT. mOTUs of the recipient post-FMT were highly correlated with mOTUs of the recipient pre-FMT (median cosine similarity of UC patient mOTUs = 0.93, CD patients = 0.95). More importantly, the results showed that mOTUs of post-FMT recipients had high similarity to mOTUs of their donors (median cosine similarity of UC patient mOTUs =0.95, that of CD patients = 0.91) (Figure 3B).

**Figure 3.**
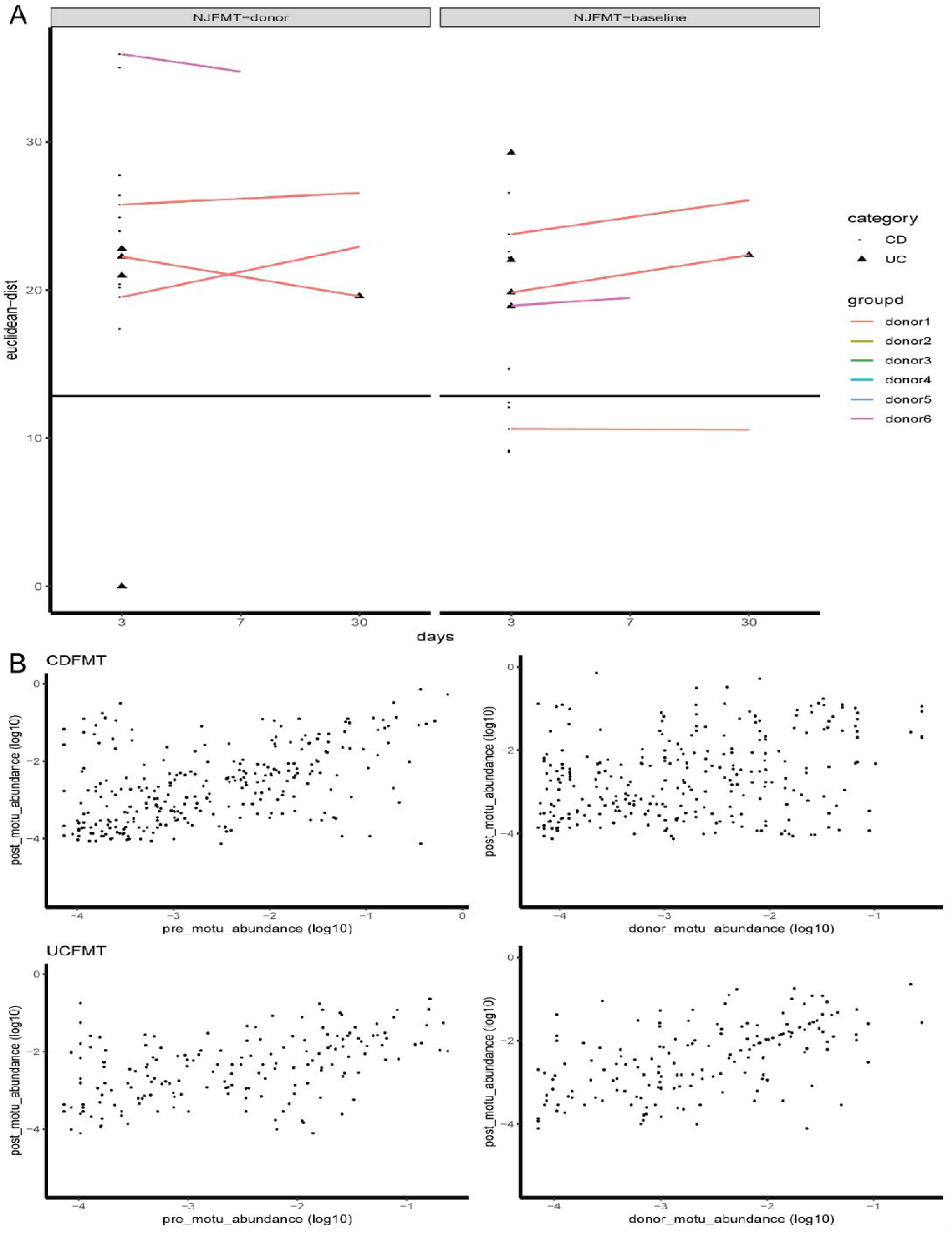
**High compositional resemblance of the gut microbiomes of post-FMT recipients and their pre-status, as well as post-FMT recipients and their donors.** (A) After FMT, the microbiota composition of most patients is further from their initial status than natural shift observed in placebo (solid black line). Additionally, recipients with the same donor (lines of the same color) may vary in their shifting tendency. (B) High consistency (median cosine similarity > 0.9) is found between post-FMT IBD patients (3 days after treatment) with their pre-FMT status, as well as with their donors.

### Bacterial engraftment at the strain level

To investigate the extent of strain level changes in our study groups, we monitored SNVs identified at baseline over all available time points. Higher levels of single-site allelic variations were observed in UC FMT recipients and CD FMT recipients compared to autologous FMT recipients from a previous paper (*21*) (P= 0.0056 and 0.148, respectively). Moreover, SNVs were found to be higher in UC FMT recipients than in CD FMT recipients (P = 0.070) (Figure 4).

**Figure 4.**
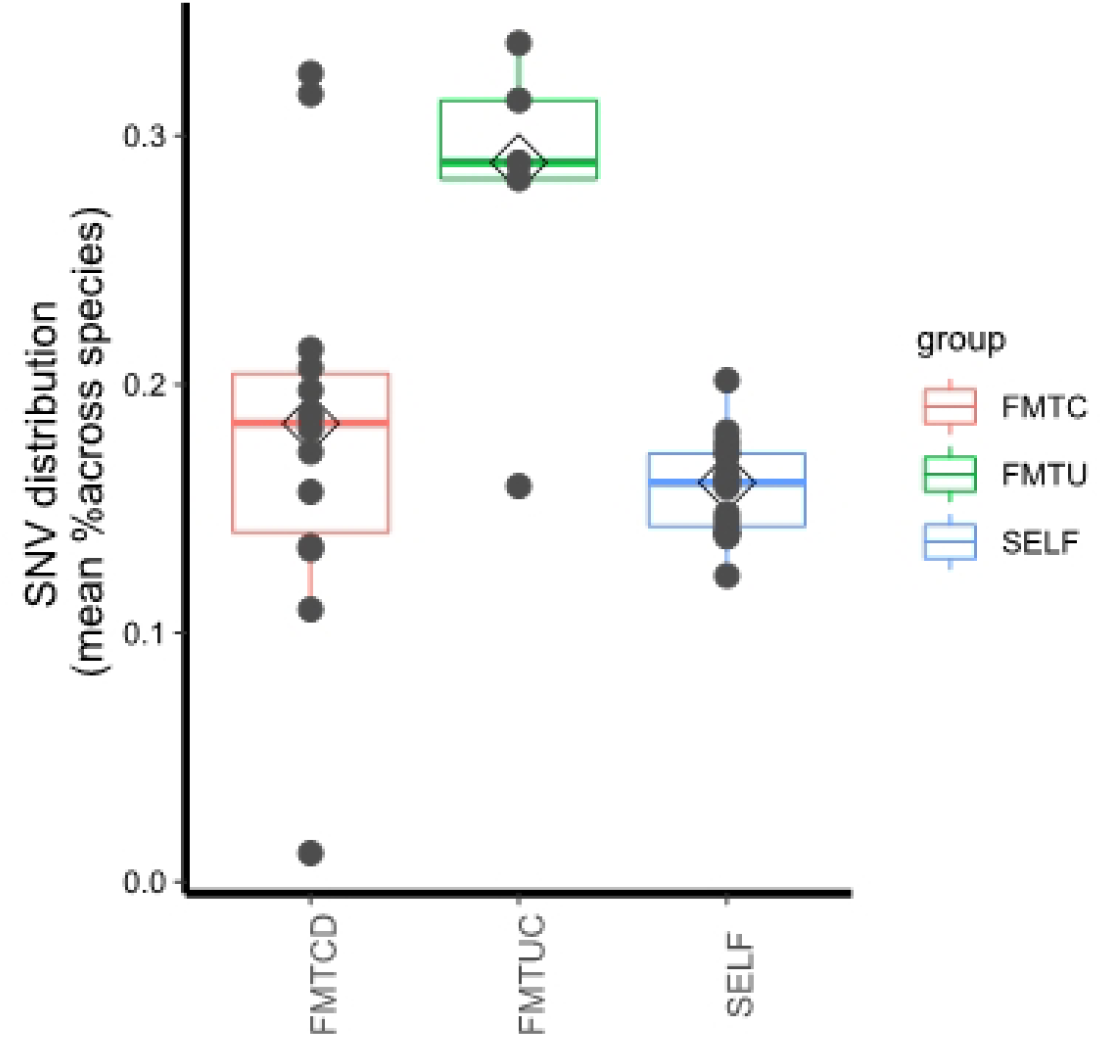
**UC recipients display higher strain level variations than CD recipients 3 days after FMT treatment.** Single-site allelic variations of UC and CD recipients after FMT treatment are a bit higher than autologous FMT recipients (p-value = 0.148 and 0.234, respectively). Single-site allelic variations of UC recipients are significantly higher than CD recipients after FMT treatment (p-value = 0.00056).

To investigate whether this increased variation was due to the transfer and establishment of donor microbiota, we followed methods described in a previously published paper (*28*), defining a set of determinant genomic positions (containing both donor- and recipient-specific SNVs) and monitoring them over time (Figure 5). For the credibility of SNVs detection, we chose species with sufficient abundance that were consistently detected in at least one donor-recipient pair. Donor-specific SNVs were most highly retained 3 days after FMT (UC: 62.8±25.3% of determinant positions across recipients, CD: 11.4±10.3%) and were still present 1 month later (UC: 46.9%, CD: 19.99 ±10.1%). This was in contrast with the much lower rates of variation observed at equivalent time points in autologous FMT recipients (9.5 ± 1.8%) (Figure S1), showing that the increased variations of gut microbiota in post-FMT patients could be attributed to donor strain transfer instead of temporal variability.

**Figure 5.**
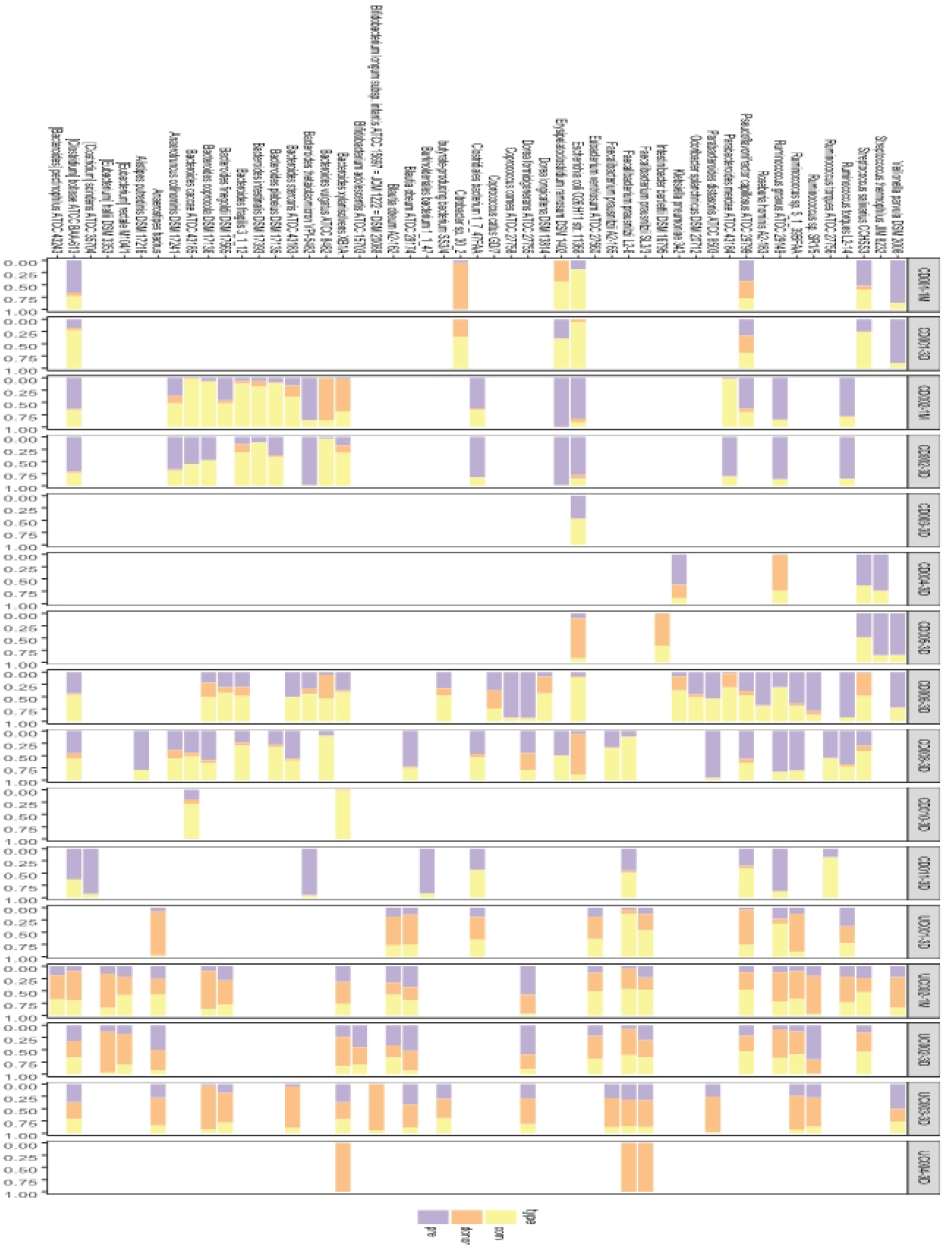
**Some donor-specific strains undergo transfer, and the existence of donor strains are highest 3 days after FMT.** The rate of donor strain transfer is greatest in recipients 3 days after FMT (UC: 62.8 ±25.3%, CD: 11.4 ±10.3%), and a portion of them persists in recipients 1 month later (UC: 46.9%, CD: 19.99 ±10.1%). Proportions of donor- and recipient-specific strains across 50 species are shown in orange and purple, respectively.

Furthermore, marked differences in colonization success were observed between UC and CD recipients who shared a donor (subjects CD-1,2,3,8, and UC-1,2). 3 days after treatment, UC-1,2 retained a higher amount of donor-specific SNVs compared to CD-1,2,3,8 (48.9%, 44.4%, 11.9%, 3.4%, 1.5%, and 9.3%, respectively). Extensive coexistence of donor and recipient strains (CD: in 44.1 ± 17.1% of shared species, UC: 21.3 ± 14.1%) was found in all other recipients, and persisted for at least one month. This suggests that novel strains can colonize the gut without replacing the indigenous strain population of the recipient. It appeared that introduced strains were more likely to be established in a new environment if the species was already present, and a pattern of donor strains establishing alongside indigenous strains of the recipient was observed. While the phenomenon of donor strain establishment occurred in both CD and UC recipients, UC patients were more susceptible to external sources of microbiota (Figure 6).

**Figure 6.**
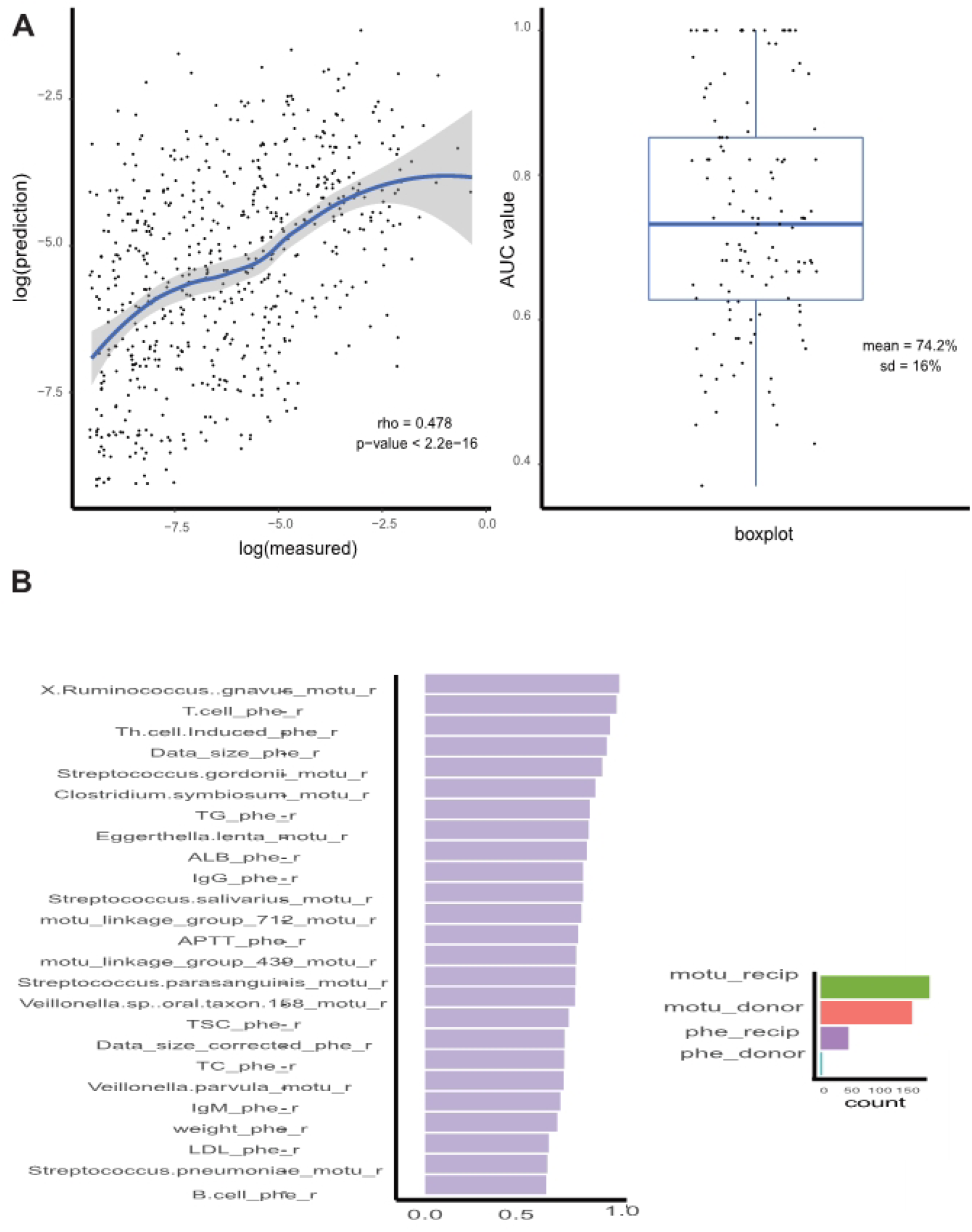
**Random forest models have the ability to predict the gut microbiota composition of post-FMT patients.** (A) Left panel shows the classification result: predicted values have a moderate consistency with true values (rho = 0.478 and p-value < 2.2e-16). Right panel shows the regression result: a boxplot of all the AUC values of each mOTU in post-FMT recipients (median AUC value = 74.2%, SD = 16%). (B) Important variables are computed across those models, defined as those with an AUC value greater than 0.90. Important variables are divided into different categories (represented by different colors). The top 25 variables are classified as the clinical parameters of recipients.

Donor strains showed different transferability under different disease status. Donor-specific strains like *Ruminococcus torques ATCC 27756, Ordoribacter splanchinicus DSM 20712, Klebsiella pneumoniae 342, Intestinaibacter bartlettii DMS 16795, Escherichia coli O26:H11 str. 11368,* and *Erysipelatoclostridium ramosum DSM 1402* only exerted strain displacement in CD patients, while donor-specific strains like *Faecalibacterium prausnitzii SL3/3, Eubacterium ventriosum ATCC 27560, Blautia obeum A2-162, Bifidobacterium longum subsp.infantis ATCC 15697 = JCM 1222 = DSM 20088, Anaerostispes hadrus,* and *Eubacterium rectale M104/1* only exerted strain displacement in UC patients (Figure 5).

### Construction of a prediction model for gut microbiota composition of post-FMT patients

According to what we have discovered in previous species-level analysis, microbiota of post-FMT recipients are a complex mixture of species from the donor, species from the recipient, and species gained from the environment. We speculated that after accounting for the gut microbiota composition of pre-FMT recipients and donors, along with the corresponding clinical metadata of the recipients, we might be able to predict the post-FMT gut microbiota of the recipients. We, therefore, performed random forest classification and regression analysis, which is non-linear and can accept categorical and continuous predictors simultaneously from our data (*25*).

To investigate whether species compositions of post-FMT patients—that is, the mOTUs profiles—were predictable, we first examined the presence of each mOTU across post-FMT recipients using the randomForest classification model, and computed the average area under the curve (AUC) (mean = 74.2%, SD = 16%). We then utilized a randomForest regression model to test the predictability of abundance of each mOTU (rho = 0.478, P < 2.2e-16). Results indicated that the presence of most (>80%) species of post-FMT recipients was highly predictable (AUC > 85%), while a small portion of species was not. The abundance of mOTUs of post-FMT recipients was moderately predictable (Figure 7A). Our results were poorer than a similar study conducted by Christopher *S. Smillie* et al. (*20*) on 19 R-CDI patients. One possible explanation for this discrepancy may be that they included other predictors in their model construction in addition to the ones we used: taxonomy, abundance, clinical metadata, sequencing depth, genome statistics, physiology, and resource utilization.

**Figure 7.**
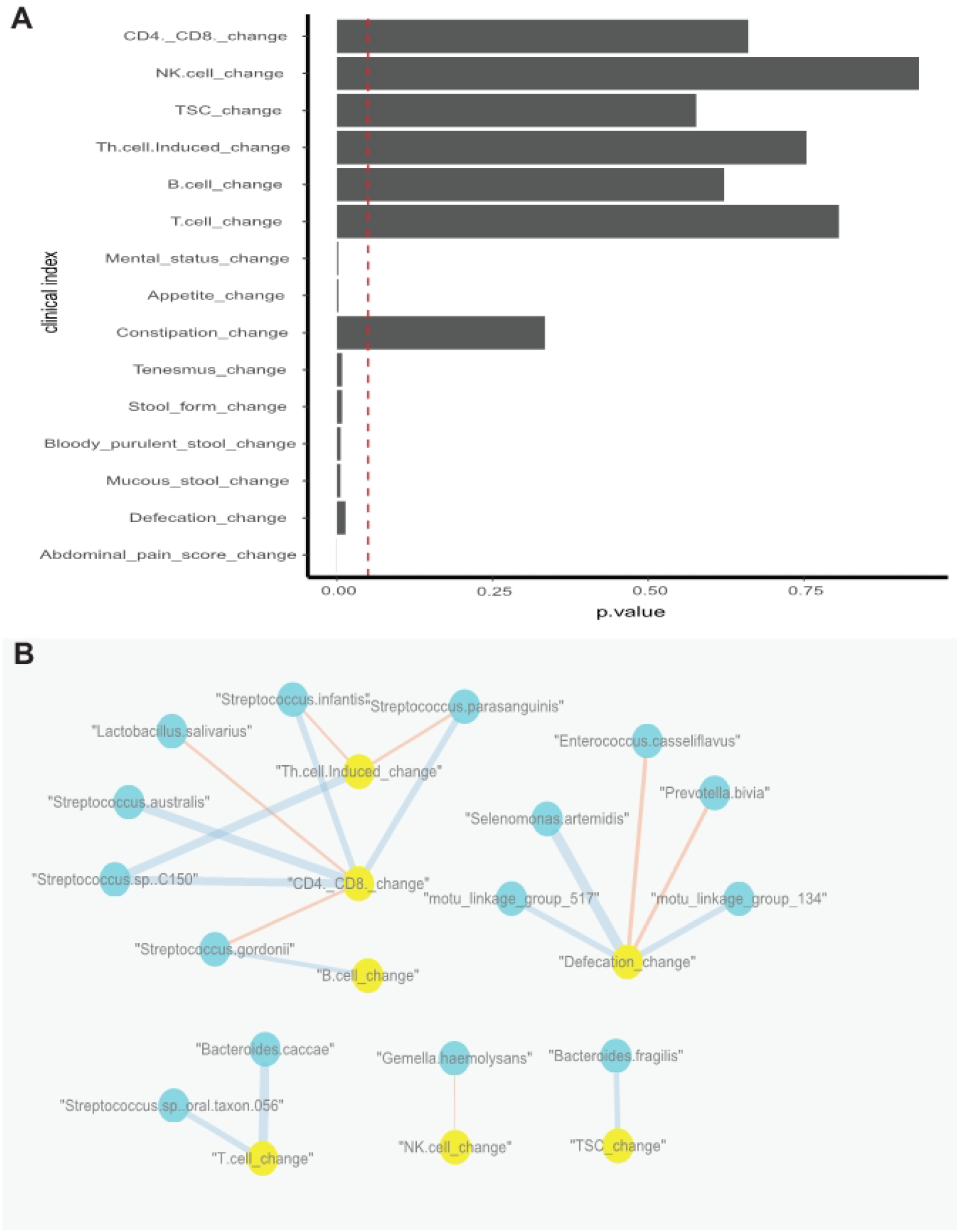
**Some clinical indexes of IBD recipients have significantly changed 3 days after FMT, and several clinical indexes correlated with changes in the mOTUs profiles of recipients.** (A) Mental status, appetite, tenesmus, etc. significantly changed 3 days after FMT (p-value < 0.05). Vertical dotted line indicates a p value of 0.05. (B) Defecation changes and CD4+CD8+ changes have relationships with several mOTUs. Blue represents a significant positive correlation, while red indicates a significant negative correlation (p-value < 0.01). The width of the lines indicates the weight of correlation.

The RandomForest model also provided an algorithm to rank the contribution of each predictor based on variable importance score. According to our analysis, among the top 40 most important variables (see Materials & Methods), the IgA score, T-cell, and Th cell-induced of the recipients were the top three clinical-related elements. *Streptococcus.anginosus, Bacteroides.plebeius, Clostridium.bolteae, Streptococcus.thermophilus*, and *X.Ruminococcus.gnavus* were the top five species in the classification model (Figure 7B). In terms of species-related factors, *Streptococcus.anginosus* was reported to be associated with colorectal cancer and *Ruminococcus.gnavus* was found to be linked with a certain type of immunological rejection.

### Clinical outcomes

Out of all 15 patients, 8 out of 11 CD patients and 3 out of 4 UC patients were relieved 3 days after FMT treatment. Clinical improvement was defined as a decrease in the Harvey-Bradshaw Index > 3 for CD, and a decrease in the Mayo score > 3 for UC (Table S1).

### Relationship between changes in clinical index and changes in gut microbiota

Potential antigens in the microflora could have pro- or anti-inflammatory effects, and it could be argued that by reacting to these antigens, an organism is mounting an autoimmune response; by extension, the chronic mucosal inflammation of IBD could be thought of as an autoimmune disease. Given this perspective, it would make sense to relate the change in clinical parameters to the abundance change of gut microbiota in response to FMT treatment. We established the relationships between the change in clinical indexes as well as in mOTUs of recipients using Spearman’s correlation. We found that defecation changes were significantly positively correlated with *Selenomonas.artemidis* and two unclassified species, and negatively correlated with *Enterococcus.casseliflavus* and *Prevotella.bivia*. Changes in CD4+CD8+, which have been identified to be higher in IBD patients than in normal people in previous studies, were significantly positively correlated with *Streptococcus.sp..C150*, *Streptococcus.infantis*, *Streptococcus.parasanguinis*, and *Streptococcus.australis*, ande negatively correlated with changes in *Streptococcus.gordonii* and *Lactobacillus.salivarius*, a probiotic bacterium that lives in the gastrointestinal tract and has a range of therapeutic properties including suppression of pathogenic bacteria (*29*). Changes in TSC were significantly positively correlated with changes in *Bacteroides.fragilis*, which is found in most anaerobic infections and can promote the induction of type 1 T helper (TH1) cells, suppress IL-17 production, and improve experimental colitis (*30*). Additionally, we tested whether the physical characteristics of patients, such as BMI, age, and disease duration, (Table S2) could affect clinical outcomes. Changes in CD4+CD8+, Th.cell.Induced (counted by Flow cytometry), and abdominal pain score were found to significantly negatively correlate with the disease start age of patients (p < 0.05), which could reflect disease duration. In addition, changes in CD4+CD8+ and Th.cell.Induced were significantly negatively correlated with the age of patients (when patients received FMT treatment). Disease duration and the age of patients were also discovered to be important features in the random forest classification model. As a result, we speculated that disease duration and age could be used as stratifying factors for IBD patients in future therapy plans (Figure 7B).

## DISCUSSION

Consistent with previous findings, our study found reduced bacterial diversity in CD and UC patients. Strain level analysis monitored across samples revealed that 3 days after FMT treatment, a certain amount of species had noticeable strain replacements. Moreover, donor-specific strains belonging to different species demonstrated differentially competitive advantages during the process of displacement, measured by their relative abundance in recipients after FMT. We also observed that same-donor recipients undergo varying degrees of gut microbiome shifts, implying that the FMT treatment effect may be patient-specific, and raising the possibility of patient stratification in clinical application.

We also aimed to identify factors that could contribute to the accurate prediction of post-FMT gut microbiota composition of the recipients. The moderate predictability of the classification and regression model suggests that the gut microbiota composition of post-FMT recipients can be recognized not through sequencing methods but through algorithms, indicating a promising future towards FMT precision treatment. In our model, we only take the species composition of the donor and pre-FMT patient, along with clinical indexes of the pre-FMT patient as predictors. There is space left to enhance the resolution of prediction accuracy. Based on previous studies concerning the etiology of IBD, factors like genetic background, non-bacterial components (virome, fungi), metabolites profile, and dietary records have the potential to account for the unexplainable part of our model.

Associations between immunological factors and clinical outcomes provide us with some limited but intriguing perspectives. CD4+CD8+, TSC, and Th.cell.Induced have been found to be associated with certain bacterial species, implying that bacteria have the potential to affect the adaptive immunity of patients. However, there are many intermediate issues to be dealt with before making a cohesive interpretation of this assumption. Combining the information from functional metagenomes and metabolomics will minimize the gap between gut microbiota and immunological responses of the recipients.

The present attractive clinical findings are mainly based on our one-hour FMT protocol for providing fresh FMT, which means the time from defecation of stool to deliver purified microbiota to patient’s intestine within one hour (*31,32*). Another factor contributing to this positive clinical response, according to our experience, might be the criteria of donor screening which is based on young age population, generally cover children and college students under 24-year-old (*33*). However, the small sample size of our study and the incomplete follow-up visits inevitably limited the scope of our results. Future studies need to include a larger study cohort and longer tracking times for the explicit identification of specific bacterial strains that may play a role in FMT treatment efficacy, and to uncover a comprehensive principle of strain displacement.

## Contributors

Conceptualization, Methodology, and Writing – Zhuye Jie, Manli Zou, Bota Cui and Faming Zhang; Revision & Editing – Faming Zhang, Huijue Jia; Acquisition of data – Bota Cui, Honggang Wang, Qiang Feng; Materia Support – Yuanqiang Zou, Xiuqing Zhang, Huanming Yang, Jian Wang; Software and Formal Analysis, Manli Zou and Zhuye Jie; Writing – Original Draft, Manli Zou, and Zhuye Jie; Supervision, Faming Zhang and Huijue Jia.

## Disclaimer

The authors declare that they have no competing interests and there is nothing to disclose.

## Patient consent

Obtained

## Ethics approval

This study was approved by BGI-IRB (BGI-R004-05)

## Availability of supporting data

Datasets are in a publicly accessible repository: The quality-controlled sequencing reads are available in the CNGB Nucleotide Sequence Archive (CNSA: https://db.cngb.org/cnsa; accession number CNP0000134)

## Additional files

Tables S1; Table S2

**Figure S1. The amount of donor-specific species gain after FMT differs, even for same-donor recipients.**

Recipients that share a donor are colored the same.

**Figure S2. A certain number of donor species display apparent transfer after FMT treatment in IBD patients.**

Heatmap and hierarchical clustering of mOTU profiles for all samples. Pre- and post-FMT CD recipients, pre- and post-FMT UC recipients, and healthy controls are separated by space.

